# SoEM-Web: a user-friendly platform for the analysis and visualization of small-organelle-enriched metagenomics data

**DOI:** 10.1101/2025.02.12.637736

**Authors:** Jooseong Oh, Hanjin Kim, Chungoo Park

**Affiliations:** School of Biological Sciences and Technology, Chonnam National University, Gwangju, 61186, Republic of Korea; Institute of Systems Biology & Life Science Informatics, Chonnam National University, Gwangju, 61186, Republic of Korea

**Keywords:** small-organelle-enriched metagenomics, marine microeukaryote diversity, web-based bioinformatics tool, environmental DNA, biodiversity assessment

## Abstract

**Background:** Marine microeukaryotes play a crucial role in global biogeochemical cycles and ecosystem functioning. However, their diversity and distribution patterns remain poorly understood. To improve the biodiversity assessment of environmental samples, we recently developed a PCR-free, small-organelle-enriched metagenomics (SoEM) method, accompanied by a bioinformatics workflow for analyzing SoEM metagenomic data. However, the complexity of these analyses typically requires extensive bioinformatics expertise, potentially limiting their broader adoption.

**Methods:** To address this challenge, we present SoEM-Web, a user-friendly web application for the interactive analysis, classification, and visualization of SoEM metagenomic data. SoEM-Web integrates all necessary procedures, including raw sequencing data preprocessing, paired-end read merging, automated taxonomic identification, and diversity visualization, into a streamlined workflow accessible through an intuitive interface. This application is designed to facilitate SoEM data analysis for researchers who may not have advanced bioinformatics skills.

**Results:** We demonstrated the capabilities of SoEM-Web by reanalyzing previously published datasets. This reanalysis not only confirmed the utility of the platform but also revealed additional taxa, emphasizing the value of updated reference datasets. The application successfully simplifies complex bioinformatics processes, enhancing accessibility and reproducibility in marine microeukaryote diversity research. SoEM-Web aims to enhance accessibility and reproducibility in marine microeukaryote diversity research. The web server is freely available at http://compsysbio.re.kr/soem-web/.

## Introduction

The rapid advancement of high-throughput sequencing technology and bioinformatics methods has propelled environmental DNA (eDNA) metabarcoding to the forefront of marine microeukaryotic biodiversity monitoring (Balasubramanian et al. 2021; Brandt et al. 2020; Brannock et al. 2016). Despite its promise, the systematic use of eDNA metabarcoding in marine biodiversity monitoring encounters several challenges (Pawlowski et al. 2022; Sigsgaard et al. 2020; Takahashi et al. 2023). These include polymerase chain reaction (PCR) bias—where different species are amplified with varying efficiencies—the need for suitable eDNA-based metabarcoding markers; and the development of efficient protocols for eDNA filtration, extraction, inhibitor removal, and amplification.

To address these limitations, researchers have developed alternative approaches such as metagenome skimming (Dodsworth 2015; Greshake et al. 2016; Linard et al. 2015), mitochondrial metagenomics (Crampton-Platt et al. 2015; Crampton-Platt et al. 2016; Tang et al. 2014), mitochondrial capture microarray (Liu et al. 2016), and mitochondrial enrichment through differential centrifugation (Zhou et al. 2013). Building on these advances, we recently developed a PCR-free small-organelle-enriched metagenomics (SoEM) method, which demonstrated superior performance in marine species identification compared to multi-marker eDNA metabarcoding (Jo et al. 2019). We subsequently refined this approach to enable efficient eDNA extraction from small-volume water samples using optimized cell disruption and DNA extraction methods (Jin et al. 2023).

Complementing these experimental advancements, we developed a bioinformatics pipeline for SoEM metagenomic analysis. This pipeline encompasses raw sequencing data preprocessing, paired-end (PE) read merging, taxonomic identification, and visualization of taxonomic diversity. However, the implementation of these *in silico* analyses typically requires expertise in various software programs, potentially posing a challenge for researchers primarily focused on wet laboratory experiments.

To bridge this gap, we developed SoEM-Web, a web-based tool that makes SoEM bioinformatics analyses accessible to users without extensive programming knowledge. SoEM-Web offers an interactive, point-and-click interface that allows users to easily specify parameters, run tools, and view metadata. The tool is freely available as an online web service at http://compsysbio.re.kr/soem-web/.

By simplifying the complex bioinformatics processes associated with SoEM analysis, SoEM-Web aims to facilitate the broader adoption of this method in marine biodiversity research. We anticipate that this tool will serve not only as a practical application for professional and research purposes but also as a valuable resource for training and educating novice marine biologists in metagenomic analysis techniques.

## Materials & Methods

The SoEM-Web platform integrates five core modules, each designed to perform specific functions in the analysis pipeline: raw sequencing data upload and preprocessing, PE read merging, taxonomic identification assignment, and taxonomic diversity visualization (**Fig. 1** and **Table 1**). The following sections provide a detailed description of each module’s functionality and implementation.

**Table 1:**
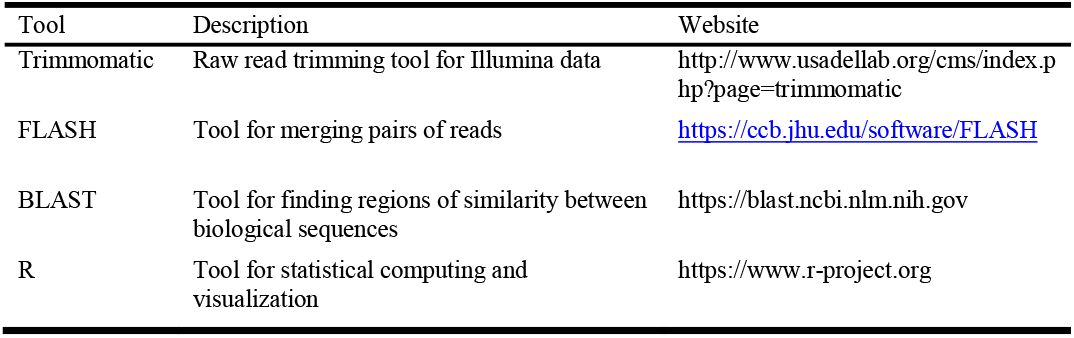
Tools for metagenomic analysis in SoEM-Web.

**Fig. 1.**
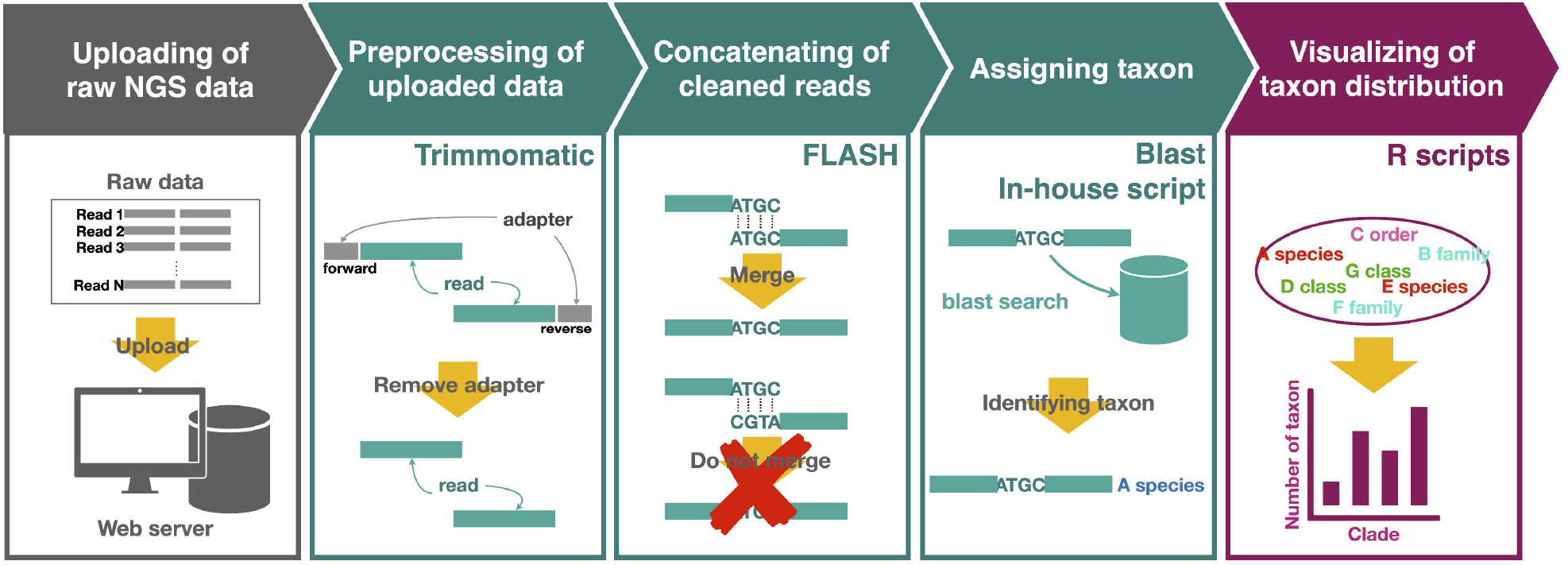
Workflow for SoEM-Web.

### Uploading and preprocessing raw next-generation sequencing data

SoEM-Web is designed to process user-uploaded FASTQ files containing DNA sequence reads. In the FASTQ format, each read is represented by four lines: an identifier beginning with “@,” the nucleotide sequence, a “+” delimiter, and corresponding quality scores (Cock et al. 2010). For optimal performance, we recommend that users input PE reads, as these can be partially overlapped to generate longer single reads, thereby enhancing the accuracy of subsequent analyses.

To optimize data handling and processing efficiency, SoEM-Web supports gz-compressed FASTQ files (e.g., fastq.gz) as input. In addition, the platform features a user-friendly drag-and-drop interface, allowing for the simultaneous upload of multiple files (Afgan et al. 2018). This feature significantly streamlines the data input process, particularly when dealing with large datasets or multiple samples.

Upon successful upload, the raw sequences undergo preprocessing using Trimmomatic v0.39 (Bolger et al. 2014). This crucial step uses default settings to remove adapter sequences introduced during library preparation and to eliminate low-quality reads.

### Merging PE reads

Following preprocessing, SoEM-Web merges cleaned forward and reverse reads that overlap to generate longer DNA sequences, a critical step for improving the accuracy of taxonomic assignments. This process is efficiently executed using FLASH (Fast Length Adjustment of SHort reads) version 1.2.11.4 (Magoc & Salzberg 2011). The FLASH algorithm is applied with carefully optimized parameters to balance sensitivity and specificity in read merging. These parameters include a minimum overlap length of 10 base pairs (bps), an average read length of 300 bps, an expected fragment length of 550 bps, and a standard deviation of fragment length set to 55 bps. The resulting merged reads form the basis for subsequent analyses, whereas unmerged reads are excluded to maintain data quality and consistency.

### Assigning taxonomic identification to merged reads

In the taxonomic identification phase, merged reads are treated as operational taxonomic units (OTUs) and mapped against four comprehensive reference databases compiled from NCBI (**Table 2**). These databases include the non-redundant nucleotide database (NT), primary marker sequences from GenBank (PM), the mitochondrial whole-genome database (MT), and the plastid whole-genome database (PT).

**Table 2:** Four reference databases available in SoEM-Web.

| Database | Abbreviation | Description |
| --- | --- | --- |
| Non-redundant nucleotide database | NT | Non-redundant nucleotide database in the NCBI nucleotide database<br>(https://www.ncbi.nlm.nih.gov/nucleotide/) |
| Primary-marker database | PM | Manually downloaded sequences from the NCBI nucleotide database through keyword searches using “*COI*,” “*COI-5P*,” “*rbcL*,” or “*tufA*” |
| Mitochondria whole-genome database | MT | All sequences of mitochondrial genomes registered in the NCBI organelle genome resource<br>(https://www.ncbi.nlm.nih.gov/genome/organelle/) |
| Plastid whole-genome database | PT | All sequences of plastid genomes registered in the NCBI organelle genome resources<br>(https://www.ncbi.nlm.nih.gov/genome/organelle/) |

The mapping process uses BLAST version 2.16.0+ (Camacho et al. 2009) with stringent parameters to ensure high-confidence matches. Specifically, an E-value threshold of 1e-10 is employed to minimize false positive alignments, and the “max_target_seqs” parameter is set to 1 to retain only the best match for each query sequence.

Taxonomic assignment is performed using the NCBI taxonomy database (Schoch et al. 2020), which provides a standardized and regularly updated classification system. Each OTU sequence is assigned the best taxonomy based on its BLAST search result and corresponding accession number. To facilitate this process, we have constructed a custom SoEM-Web database that integrates NCBI accession numbers, taxonomic IDs (taxids), taxonomy nodes, and hierarchical information. This custom database is built using taxdump files with trackable taxids (https://ftp.ncbi.nih.gov/pub/taxonomy/taxdump_archive/) and accession2taxid mapping files from the NCBI repository. To ensure the most up-to-date taxonomic information, the four reference databases (NT, PM, MT, and PT) and taxonomy archive files undergo biannual updates.

### Visualizing taxonomic diversity and distribution patterns

The final module of SoEM-Web focuses on the visual representation of taxonomic composition and diversity. We have implemented a robust visualization module using R v4.4.2 and the ggplot v3.5.1 library (Wickham 2016), complemented by custom in-house scripts. All generated graphs can be easily downloaded in multiple formats (.pdf, .jpeg, or .tiff), making them convenient for inclusion in publications, presentations, or further analyses.

### SoEM-Web analysis protocol

The SoEM-Web analysis protocol consists of six main steps, each designed to guide users through the process of analyzing metagenomic data. Below is a detailed description of each step:

Step 1: Data Upload

1. Access the SoEM-Web site at http://compsysbio.re.kr/soem-web/.
2. Click “Upload” in the Tools menu on the left panel.
3. Drag and drop FASTQ files into the middle panel, or click “Choose local files” to select files.
4. Click “Start” to initiate the upload. Uploaded datasets will appear in the History panel.

Step 2: Quality Control

5. Click “Trimmomatic” in the Tools menu.
6. Select “Paired-end (two separate input files)” for PE data.
7. Choose the appropriate input FASTQ files for R1 and R2.
8. Click “Run Tool” to start the quality control process.

Step 3: Read Merging

9. Click “FLASH” in the Tools menu.
10. Select “Individual datasets” in the first drop box.
11. Choose the appropriate forward and reverse read datasets.
12. Set the parameters: minimum overlap (default: 10), average read length (default: 300), fragment length (default: 550), fragment length standard deviation (default: 55), and minimum fragment size (default: 400).
13. Click “Run Tool” to merge the reads.

Step 4: Taxonomic Annotation

14. Click “NCBI BLAST+” in the Tools menu.
15. Select the merged reads dataset.
16. Choose the desired reference database (NT, PM, MT, or PT).
17. Set the E-value (default: 1e-10) and select the output format (default: Tabular).
18. Specify the minimum alignment length (default: 400).
19. Click “Run Tool” to perform the BLAST search. Step

Step 5: Taxonomic Assignment

20. Click “Assign Taxa” in the Tools menu.
21. Select the BLAST output dataset.
22. Click “Run Tool” to assign taxonomic names to the merged reads.

Step 6: Visualization

22. Click “Visualization” in the Tools menu.
23. Select the taxonomic assignment output dataset.
24. Enter the sample name, plot title, and axis labels, and specify plot dimensions.
25. Choose the desired output file type (.pdf, .jpeg, or .tiff).
26. Click “Run Tool” to generate the visualization.

Throughout the analysis, users can monitor job progress and access intermediate results via the History panel. This feature allows users to rerun analyses with modified parameters if necessary, enhancing the flexibility and iterative nature of the analytical process.

## Results

### Implementation of SoEM-Web

We developed SoEM-Web, a user-friendly web-based application for analyzing and visualizing PCR-free, small organelle-enriched metagenomics data, using the Galaxy platform (Afgan et al., 2018b). The application integrates open-source bioinformatics tools and custom Python and R scripts to provide a comprehensive analysis pipeline. The SoEM-Web interface consists of three main components (**Fig. 2**): the Tools menu, the Interface panel, and the History panel. These components work together to guide users through the analysis process, which includes six main steps: data upload, quality control, read merging, taxonomic annotation, taxonomic assignment, and visualization. The detailed protocol for each step is described in the Methods section.

**Fig. 2.**
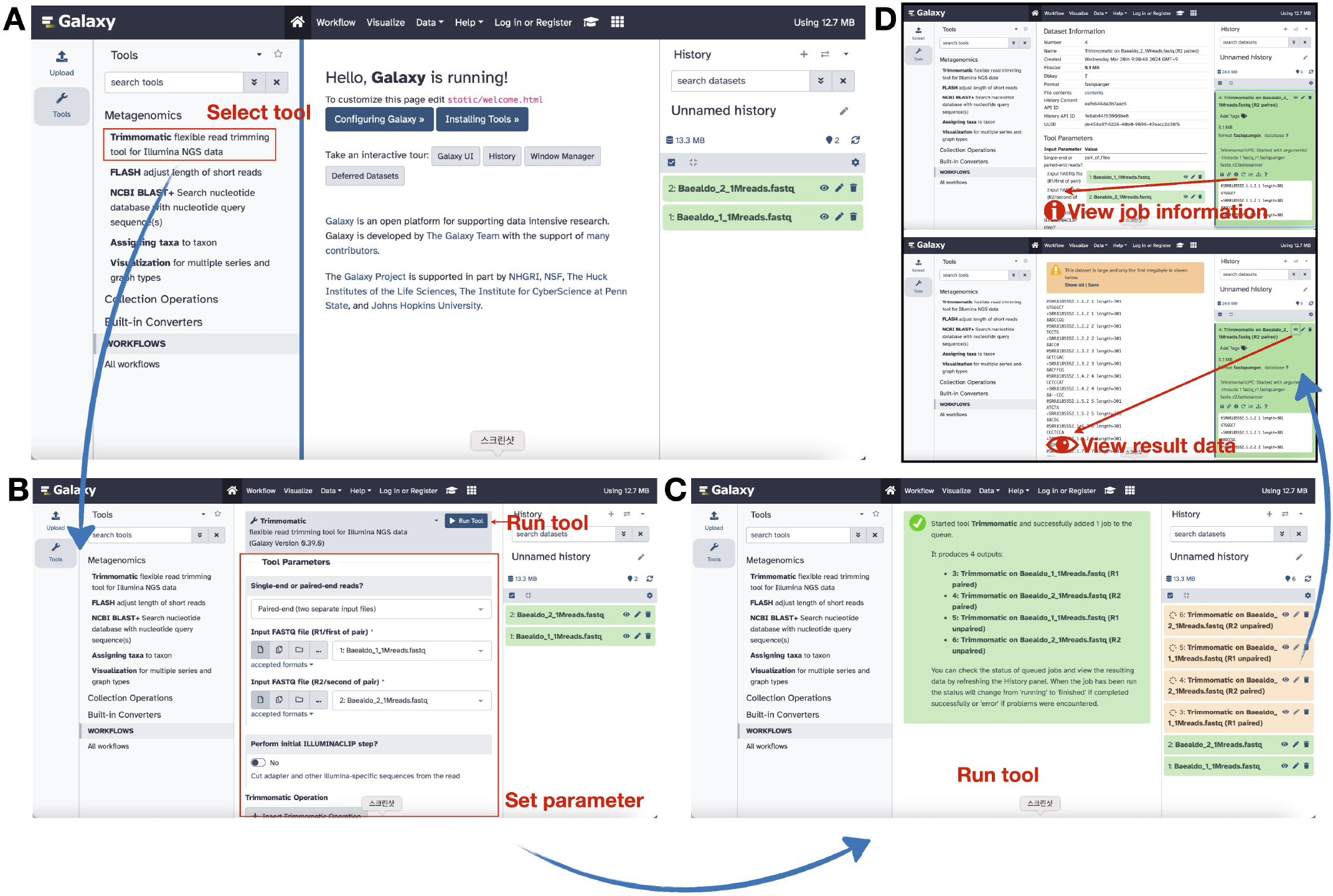
SoEM-Web interface. The SoEM-Web analysis interface consists of three main panels: the “Tools” section on the left, the “Main” panel in the center, and the “History” panel on the right. (A) The Tools section lists the available SoEM data analysis tools, allowing users to select the tool of interest. (B) After selecting the tool, users can configure input parameters and execute the tool. (C) Using the “Run” tool, users can monitor the progress of the job. (D) The History panel displays all outcomes and information about the tools used.

### Case study analysis

To demonstrate the functionality of SoEM-Web, we reanalyzed two metagenomic sequence datasets from a previous SoEM assay (Jo et al. 2019). The samples, Baealdo and Yamido, were originally sequenced using Illumina MiSeq (v2, 301-cycle). Following the SoEM-Web protocol, we processed the raw data through all six steps. After quality control, 35.1 and 34.5 million reads were obtained from the Baealdo and Yamido samples, respectively. PE read merging resulted in 9.1–9.7-million-longer reads, approximately half of the input reads (**Table 3**). The BLAST search against the NCBI NT database and taxonomic assignment identified 444 and 480 OTUs at the species level for Baealdo and Yamido, respectively. Notably, compared to the original study (Jo et al. 2019), SoEM-Web identified more species: 444 vs. 372 for Baealdo and 480 vs. 441 for Yamido. Between 22% and 37% of species were uniquely detected at each site using each method (**Fig. 3**). The visualization step allowed us to generate clear and informative plots of the taxonomic diversity and distribution patterns, facilitating the interpretation of our results.

**Table 3:** Summary statistics of the case study.

| Data | Total reads<br>(raw data) | Total read base<br>pairs<br>(raw data) | Total reads<br>(trimmed data) | FLASH-assembled<br>read-pairs to<br>single-end | Assembled<br>reads (%) |
| --- | --- | --- | --- | --- | --- |
| Baealdo | 36,503,228 | 10,817,749,358 | 35,108,854 | 9,096,882 | 51.82% |
| Yamido | 36,847,740 | 10,885,397,232 | 34,539,260 | 9,739,341 | 56.40% |

**Fig. 3.**
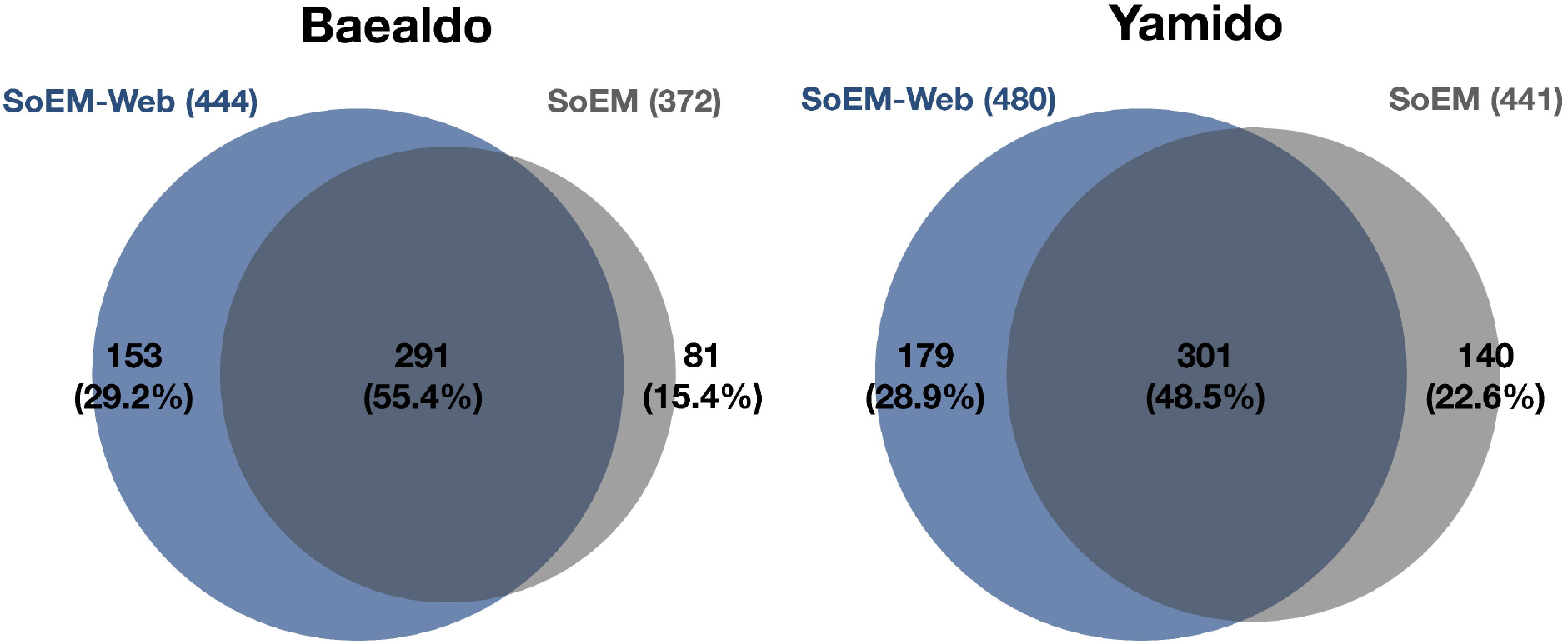
Comparison of analyzed results between SoEM-Web and those reported by Jo et al. (2019) The comparison is based on species-level information collected during the analysis.

## Discussion

The development of SoEM-Web represents a significant advancement in metagenomic analysis, particularly for studying marine microeukaryote diversity. Although the SoEM method was developed to overcome the limitations of PCR-based metabarcoding approaches, analyzing SoEM data still required substantial bioinformatics expertise. SoEM-Web addresses this challenge by providing a user-friendly interface for complex metagenomic analyses, making the process more accessible to a broader range of researchers.

SoEM-Web offers several key advantages. First, its user-friendly interface makes it accessible to researchers without extensive bioinformatics expertise. Second, it provides a standardized analysis pipeline, from raw sequencing data preprocessing to taxonomic assignment and visualization. Third, this standardized approach enhances reproducibility, a critical aspect of scientific research. Lastly, regular updates to reference databases ensure accurate taxonomic identification.

Our case study, which reanalyzed data from the study by Jo et al. (2019), demonstrates the efficiency and ease of use of SoEM-Web. We successfully performed the entire analysis without requiring complex bioinformatics expertise. The step-by-step protocol and user-friendly interface significantly streamlined the process, from quality control of raw sequencing data to read merging, taxonomic annotation, and final visualization. The ability to adjust parameters at each stage further highlights the flexibility of SoEM-Web.

Interestingly, our reanalysis revealed several notable differences from the original study. We identified more species in both the Baealdo (444 vs. 372) and Yamido (480 vs. 441) samples. Moreover, 22%–37% of species were uniquely detected by each method. These differences can primarily be attributed to the continuous expansion of the GenBank database, which approximately doubles in size every 18 months (Sayers et al. 2023). This expansion provides more comprehensive reference data for taxonomic assignment during reanalysis. The discrepancies in species identification also highlight the impact of changes in taxonomic naming and classification, underscoring the need for regular updates to taxonomic annotation and reference databases.

Furthermore, as pointed out by (Jin et al. 2020), the presence of erroneous taxonomic information in public databases emphasizes the importance of careful data curation. These findings demonstrate the value of regularly reanalyzing metagenomic datasets. As databases and analysis tools continue to improve, reanalysis of previously published datasets can uncover additional taxa and provide more accurate taxonomic profiles. However, users should be aware of the limitations associated with reference databases and exercise caution when interpreting results, particularly when comparing analyses conducted at different time points or with different database versions.

While SoEM-Web significantly simplifies the metagenomic analysis process, there is still room for further enhancement. Future developments could focus on incorporating more advanced visualization tools and integrating machine learning approaches to improve taxonomic classification. Additionally, continuously refining the pipeline to adapt to emerging sequencing technologies and evolving database structures will be crucial to maintaining the relevance of SoEM-Web in the field of metagenomic analysis.

## Conclusions

SoEM-Web, presented in this study, is a user-friendly web application designed to meet the growing demand for accessible metagenomic analysis tools in marine biology and oceanography. By streamlining the entire analysis process from raw data preprocessing to visualization, it empowers researchers to investigate marine microeukaryote diversity more efficiently. Our case study demonstrates the potential of SoEM-Web to uncover additional taxonomic information when reanalyzing existing datasets, underscoring its value in the context of rapidly expanding sequence databases. As metagenomics continues to evolve, SoEM-Web aims to support the marine research community in understanding microeukaryote biodiversity and its role in ocean ecosystems, while also promoting critical interpretation of results within the limitations of current technologies and taxonomic frameworks.

